# Graves’ disease-associated TSHR gene is demethylated and expressed in human regulatory T cells

**DOI:** 10.1101/2022.10.24.513489

**Authors:** Ahto Salumets, Liina Tserel, Silva Kasela, Maia Limbach, Lili Milani, Hedi Peterson, Kai Kisand, Pärt Peterson

## Abstract

Epigenetic changes at specific genetic loci and the activation of transcriptional repressor FOXP3 are needed to establish and maintain the regulatory T cell (Treg) lineage. Here we studied the DNA methylation profiles in CD4^+^ CD25^+^ Tregs and CD4^+^ CD25^−^ conventional T cells (Tconvs) from healthy individuals and identified a wide range of differentially methylated CpG sites (DMPs). Overall, Tregs had more hypomethylated DMPs and contained more CpG sites, which on the cell population level were less defined in their methylation status. We identified top hypomethylated CpGs in Tregs close to CENPM, IKZF2, and LYST and hypermethylated sites at the THEMIS, SCML4, and ADD3 genes. Among others, DMPs were enriched for the transcriptional repressor Kaiso binding motifs. Interestingly, in Tregs we found hypomethylation and increased expression of the TSHR gene, which is a risk gene for Graves’ disease (GD). However, subsequent DNA methylation profiling in healthy individuals and GD patients revealed only 19 DMPs and no change at the TSHR locus, indicating that Tregs in GD patients share a similar methylation pattern with healthy controls. Together, we show Treg-specific hypomethylation and expression of the TSHR gene, prompting additional scenarios to explain the genetic link and role of anti-TSHR autoantibodies in GD.

## Introduction

Regulatory T cells (Tregs) confer immunological self-tolerance and homeostasis via their immunosuppressive functions. Although the impact of Tregs on the development of multifactorial polygenic autoimmune diseases remains open, their depletion, or their key genes, leads to the emergence of autoimmunity [1, 2].

Treg phenotype is induced by forkhead box protein P3 (FOXP3) [3, 4], which acts as a transcriptional repressor under T cell receptor (TCR)-stimulated conditions [5, 6]. In addition to the FOXP3 expression, the Treg function is epigenetically controlled to maintain the accessibility of regulatory regions in its signature genes [7, 8]. The Treg-specific hypomethylation is cardinal to maintain functionally stable Treg lineage, while FOXP3 can be temporarily expressed after conventional T cell activation [3]. In Tregs, DNA is demethylated within the first intron of FOXP3 as well as in CTLA4, IL2RA (CD25), IKZF2, IKZF4, and TNFRSF18 (GITR) genes [7, 9] with high expression. Furthermore, genome-wide association studies with autoimmune diseases have revealed single nucleotide polymorphisms in regulatory regions of IL2RA and CTLA4, and that autoimmunity-related polymorphisms are enriched in Treg-specific demethylated regions and super-enhancer regions [10].

Like other endocrine autoimmune diseases, Graves’ disease (GD) affecting the thyroid gland, has been associated with impairment of Tregs. Several studies have reported reduced numbers of Tregs in GD patients [11–13] and a recent meta-analysis confirmed lower proportions of Tregs in untreated GD but not in untreated Hashimoto’s thyroiditis patients [14] [15]. The hallmark of Graves’ hyperthyroidism is the presence of anti-TSHR autoantibodies [16]. Thyroid stimulating hormone receptor (TSHR) is a G-protein coupled receptor and autoantibodies induce cAMP signalling pathway, leading to thyroid cell proliferation and thyroid hormone synthesis and secretion [16].

Intriguingly, TSHR has a strong connection with GD not only as of the autoantigen but also through the genetic polymorphisms in its intron 1 region [17]. So far, two mechanisms have been proposed to explain the association of TSHR polymorphisms with GD. The first, proposed by Brand et al. [18], suggested that the disease-associated SNPs regulate TSHR mRNA splicing, which would result in increased expression of a more autoantigenic TSHR subunit in the thyroid. Alternatively, Pujol-Borrell and colleagues [19, 20], proposed that the disease-associated SNPs cause a reduced TSHR expression in the thymus, leading to decreased central tolerance and increased risk of autoimmunity to TSHR. Supporting the role of thymic tolerance, Tomer and colleagues identified an open chromatin region marked by IFNα-induced H3K4me1 enrichment to overlap with GD-associated SNPs in TSHR intron 1 [21]. They showed that sustained IFNα production during viral infections in combination with the disease-associated variant in the regulatory region located in intron 1 of the TSHR gene permits inhibitory transcription factor binding that leads to reduced TSHR expression in the thymus and the escape of TSHR-reactive T cells from central tolerance.

In this study, we investigated the differential DNA methylation in human Tregs and Tconvs and identified multiple novel differentially methylated CpG sites. In a genome-wide analysis, Tregs had more CpG sites with intermediate methylation levels and enrichment for transcription factor Kaiso binding motifs. Interestingly, we identified hypomethylation of the CEP128-TSHR locus and upregulation of TSHR transcripts in Tregs. As Tregs have been implicated in GD, we studied the differential methylation in Tregs from GD and healthy controls and found few changed CpGs but no differential methylation in the TSHR gene.

## Material and Methods

### Study participants

We studied DNA methylation profiles from two groups of individuals. The first group contained paired samples of seven healthy individuals from whom Treg (CD4^+^CD25+) and Tconv (CD4^+^CD25^−^) cells were extracted. The second group consisted of 23 individuals (12 healthy and 11 GD patients) from whom only Tregs (CD4^+^CD25^+^) were obtained. Additionally, Tregs and Tconvs were extracted from 3 healthy males for gene expression analysis. All the participants were older than 18 and written informed consent to participate in the study was obtained from everyone before the recruitment. The diagnoses of GD patients were defined by a specialist endocrinologist using standard clinical and laboratory testing. The study was approved by the Ethics Review Committee of Human Research of the University of Tartu, Estonia according to permissions no 206/T-4 (August 25, 2011), no 272/T-12 (August 21, 2017), and no 275/M-17 (November 20, 2017). The experiments were performed in compliance with the Helsinki Declaration. Information about the age and sex of the study participants that remained in the analysis after data quality control is shown in **Sup. Table 10**.

### Extraction and sorting of blood cells

The peripheral blood mononuclear cells (PBMC) were extracted using Ficoll-Paque (GE Healthcare, Chicago, IL, USA) gradient centrifugation. We then purified CD4 CD25 and CD4^+^CD25^−^ T cells using CD4^+^CD25^+^ Regulatory T Cell Isolation Kit (#130-091-301) and AutoMACS technology (Miltenyi Biotec) according to the manufacturer’s protocol. The purity of extracted Tregs was determined with fluorescence conjugated antibodies against CD3, CD4 and CD25 (on a FACSCalibur flow cytometer (BD Biosciences, San Jose, CA, USA) [22]. The collected cell populations were stored as cell pellets in a −80 °C freezer.

### DNA extraction, bisulfite conversion and Illumina EPIC array

The genomic DNA was isolated from the purified cell pellets using the QIAmp DNA Mini Kit according to the manufacturer’s protocol (Qiagen, Hilden, Germany). DNA was precipitated in isopropanol, washed in 70% ethanol and resuspended in 1×TE buffer. The purity and concentration of the DNA samples were measured with the NanoDrop ND-1000 spectrophotometer (Thermo Fischer Scientific, Waltham, MA, USA), and an aliquot of 500□ng of genomic DNA was treated with sodium bisulfite using the EZ DNA Methylation Kit according to the manufacturer’s instructions (Zymo Research, Irvine, CA, USA). The Infinium Methylation EPIC BeadChip technology (Illumina, San Diego, CA, USA) was used to identify the DNA methylation levels in the samples.

### RNA extraction and gene expression measurement

RNA from sorted Treg and Tconv cells was extracted using QIAzoland miRNeasy Micro Kit combined with recommended RNase-free DNase I treatment (all from Qiagen, Hilden, Germany). The quality of extracted RNA was assessed using Bioanalyzer and RNA 6000 Nano Kit (Agilent). Gene expression levels were obtained using Human Clariom S arrays on GeneTitan MC Instrument International (Thermo Fisher Scientific).

### Data analysis

For the pre-processing and quality control of DNA methylation data, we used R package minfi (1.36.0) on original IDAT files from HiScanSQ scanner. We used “noob” method for background correction, dye-bias normalization and “functional normalisation” for between-array normalisation to remove unwanted variation by regressing out variability explained by the control probes and to preserve the global methylation differences. Subsequently, CpG sites were filtered out based on the following criteria: 1) detection p-value >0.01 in at least one sample; 2) CpGs that mapped to sex chromosomes (except for separate X chromosome analysis); 3) polymorphic CpGs (i.e. CpGs that are situated in the SNPs); 4) cross-reactive probes specific to EPIC array [23]. After the pre-processing, the total number of CpG sites for downstream analyses was 766,126 for Treg vs Tconv dataset and 726,882 for GD vs healthy control dataset. We annotated CpG sites using an annotation package IlluminaHumanMethylationEPICanno.ilm10b4.hg19. During the quality control, we removed one sample from the first analysis (one female Treg sample) and two from the second (two healthy females) due to data quality issues. Therefore, we studied the differences in DNA methylation between Tregs and Tconvs using 5 female and 1 male samples except for X chromosome analysis when only 5 females were used. Regarding to DNA methylation differences between Tregs from Graves’ patients and healthy controls, we used samples from 8 female and 3 male Graves’ patients, and 7 female and 3 male healthy controls. Similarly, for X chromosome analysis only females were used (8 Graves’ patients and 7 healthy controls).

We used R package “oligo” (version 1.54.1) to pre-process the gene expression data. More specifically, we used RMA algorithm (robust multichip average) to perform background adjustment, across-array normalisation, and summarisation. The log_2_ data was used for the quality control and analysis. We annotated probes on Human Clariom S array with an annotation file available on Affymetrix homepage. In quality control we removed one Treg sample from the analysis due to quality issues and filtered out lowly expressed genes (cut-off was set on 4). The final number of genes included into the analysis was 11,453. In total, for differential analysis, we used Tregs from 2 male and Tconvs from 3 male participants.

We used limma package (3.46.0) for differential DNA methylation and gene expression analyses. If the data originated from the same individual, we included individual as a covariate in linear model. We did all statistical tests on DNA methylation using M-values (M-value = log_2_(M (methylated intensity)/U (unmethylated intensity))). Statistical tests on gene expression were performed on log_2_ data. Throughout the analysis, we used false discovery rate (FDR) to adjust p-values. We obtained differentially methylated regions by using DMRcate R package (version 2.4.1). For enrichment analysis, we used g:Profiler [24] with default parameters. Additionally, we used custom GMT in g:Profiler to obtain enrichment results for The Molecular Signature Database (MSigDB) C7 immunologic signatures sets.

## Results

### Widespread CpG hypomethylation in Tregs

Previous studies identified a relatively low number of differentially methylated CpG sites between human Tregs and Tconvs [25–27]. We performed DNA methylation analysis of healthy human Tregs and Tconvs and found remarkably more sites when studying paired samples from the same individuals. In total, we found 16,624 autosomal CpG sites that were differentially methylated between Tregs and Tconvs with FDR ≤ 0.05 (**Fig 1A,** all significant results are shown in **Sup. Table 1**). In agreement with previous studies, hypomethylation was more prevalent among Tregs as we found 12,082 hypomethylated and 4,542 hypermethylated CpG sites in Tregs compared to Tconvs. The differentially methylated positions (DMPs) were most frequently located in gene body (41.7%, gene body and exon bands combined), however, 30% of DMPs were in or close to gene regulatory regions (TSS1500, TSS200, 5’UTR, 1st exon). Proportionally, there were more hypermethylated sites (36.5%) in the promoter area compared to hypomethylated CpG sites (27.7%) (**Fig 1B**). When we studied their location in the CpG islands, we found a majority of sites in open sea region (76.7%) although 15.8% were either in CpG islands or in shore regions (0–2 kb from islands). We found more hypermethylated DMPs in CpGs islands and shores (29%) in comparison with hypomethylated DMPs (10.8%) (**Fig 1C**).

**Fig 1.**
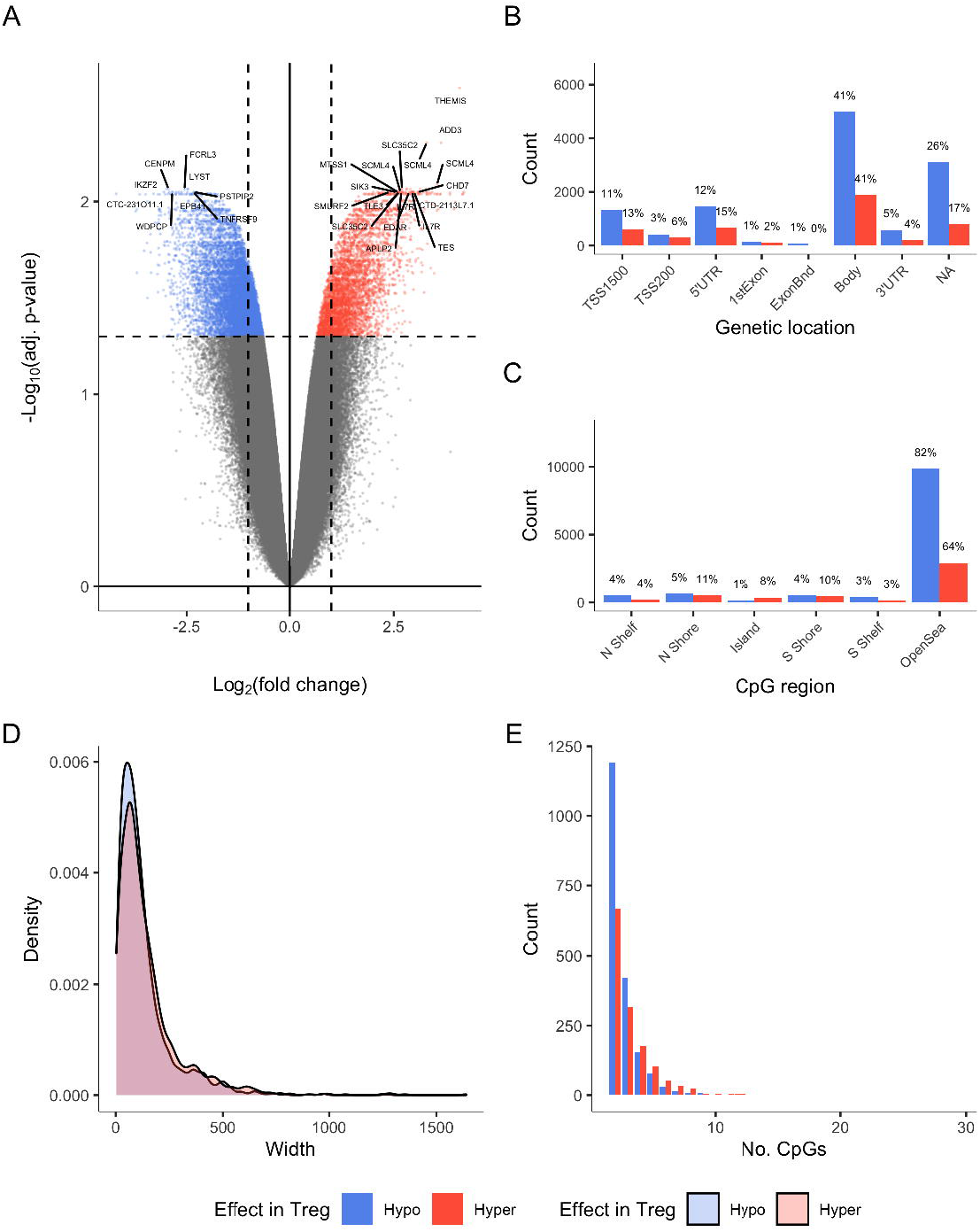
DNA methylation differences between Tregs and Tconvs. (A) Results of differential analysis represented as a volcano plot. Differentially methylated CpG sites (DMPs) are coloured red and blue corresponding to hypo- and hypermethylation in Tregs. Additionally, the genes where the top DMPs lie are highlighted. The y-axis shows −log_10_ of FDR adjusted p-values from linear regression models where individual was used as a covariate and the x-axis shows log_2_ of fold change. The number of tests was 766,126 (the number of CpGs after pre-processing) (B-C) The location of DMPs in genes and in relation to CpG islands, respectively. The DMP count is shown relative to their location and percentages indicate the proportion of hypo- and hypermethylated DMPs in a particular region. (D) Distribution of widths of hypo- and hypermethylated differentially methylated regions (DMRs) in base pairs. (E) Distribution of the number of CpGs in hypo- and hypermethylated DMRs. (A-E) Statistical analysis was done on Tregs and Tconvs from 6 individuals.

In addition to the specific CpG sites, we identified differentially methylated regions (DMRs) containing concordantly methylated and closely located CpGs (within 500 bp). We found 3304 such regions (FDR ≤ 0.05) among which 1909 were hypomethylated and 1395 were hypermethylated in Tregs (**Sup. Table 2**). The analysis of DMRs showed the hypomethylated regions to be slightly smaller in size (**Fig 1D**) and usually contained only 2 CpGs (**Fig 1E**).

We identified several differentially variable CpG positions (DVPs). These CpG sites have higher variance in their DNA methylation levels between the cell types whereas the means of the DNA methylation are not necessarily different. In total, we found 27 such sites with 24 of them showing higher variance in Tregs (**Fig S1**, **Sup. Table 3**).

### Differentially methylated CpGs reveal known and novel sites

Among DMPs in Tregs, the top hypomethylated sites were close to centromere protein CENPM, transcription factor IKZF2 (Helios), lysosomal trafficking regulator LYST, immunoglobulin receptor superfamily member FCRL3 and co-stimulatory immune checkpoint molecule TNFRSF9 (CD137/4-1BB) (**Fig S2A, Sup. Table 1**). The top hypermethylated sites were located nearby the THEMIS gene with a regulatory role in the thymic T-cell selection. The other hypermethylated CpG sites were close to SCML4, belonging to a polycomb family mediating transcriptional repression, ADD3, adducin involved in the spectrin-actin binding and controlling the actin polymerization, and GIMAP8, GTPase of the immunity-associated protein family (**Fig S2A**). Among DVPs, we found CpG sites close to ZC3H12A, which codes an endoribonuclease that degrades mRNAs involved in immune cell activation (**Fig S2B**).

We grouped the DMPs by their locations within the corresponding genes and focused on Treg signature genes. We found IKZF2, a transcription factor, which expression peaks at the Treg precursor stage, is highly hypomethylated in gene body while hypomethylation in the TIGIT gene, encoding a co-inhibitory receptor that suppresses Th1 and Th17 cells, covers the entire gene from 5’ to 3’ UTR. (**Fig 2A**). In addition, ETS1, IKZF4, FCRL3, TNFRSF9 and CTLA4 were hypomethylated in their regulatory regions while hypermethylation of these regions was characteristic for SCML4, THEMIS and NR4A3.

**Fig 2.**
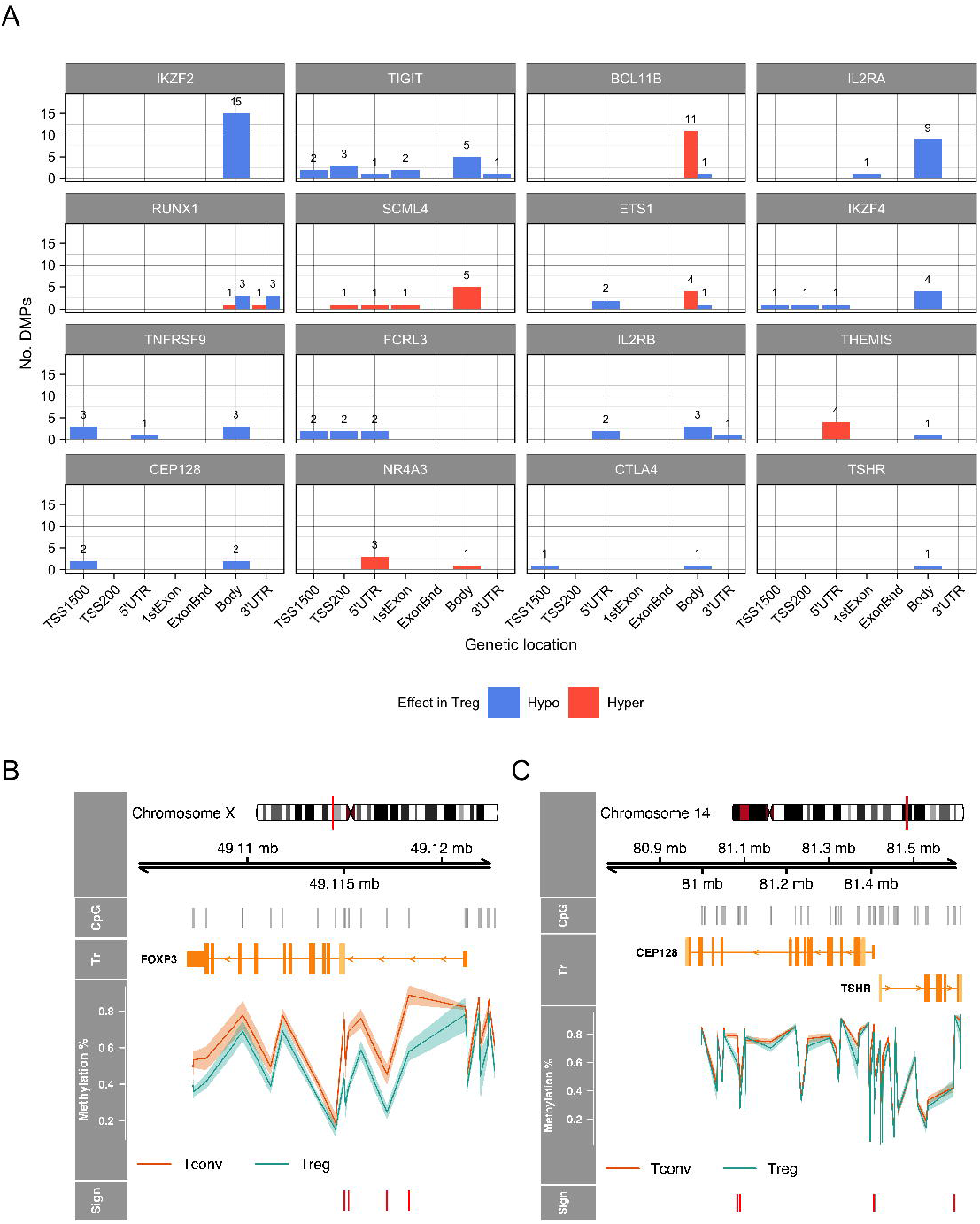
Differential methylation of specific genes in Tregs and Tconvs. (A) The number of DMPs from differential analysis (shown in Fig 1A) in specific genes with their genetic location. Blue and red colour mark hypo- and hypermethylation in Tregs, respectively. (B) Methylation levels of CpGs in FOXP3 and (C) CEP128/TSHR region with primary transcript (track denoted as “Tr”) and chromosomal information. The red stripes in “Sign” track indicate the location of DMPs. (A-C) The data consists of methylation profiles of Tregs and Tconvs from 6 individuals.

We did a separate analysis on the X chromosome using only females due to the imbalance of X-chromosome between the sexes and its inactivation (**Fig S3, Sup. Table 4-5**). We identified 28 DMPs and 6 DMRs with top DMPs and top DMR being located on the FOXP3 gene that encodes the well-known master regulator of the development and function of Tregs. Expectedly, the hypomethylation trend was present within and nearby FOXP3, particularly in intron 1, close to the CNS1 and CNS2 regions (**Fig 2B**, boxplots of individual significant sites are shown in **Fig S4A**).

Interestingly, we found that Tregs have hypomethylated CpGs within CEP128/TSHR locus on chromosome 14 (**Fig 2A, C**, significant sites individually on boxplots **Fig S4B**). The CEP128 and TSHR genes have their 5’ regulatory regions located 14 kb apart, head-to-head, and both genes have been identified as risk loci for GD [28].

### Transcriptome changes in Tregs

We next studied the Treg and Tconv transcriptome changes and used buffy coats to magnetically separate Tregs and Tconvs. We used the Affymetrix GeneTitan microarray platform and found 1552 differentially expressed genes between Tregs and Tconvs (**Sup. Table 6**). We took a closer look on top genes as well as known genes in Tregs (**Fig 3**). We found the differential expression of the TSHR gene in Tregs with its expression being nearly non-existent in Tconvs while it was lowly expressed in Tregs. Furthermore, the TSHR expression is increased specifically in the memory Treg population, albeit on a low level, but not in any other CD4^+^ T cell subtypes, according to the DICE database (**Fig S5**). Additionally, we observed higher expression of many Treg signature genes such as FOXP3, CTLA4, TIGTI, IKZF2 and NR4A1 while Tconvs showed higher expression of chemokine CXCL9, oligosaccharide maturation involved MAN1C1, ankyrin ANK3, ankyrin repeat protein ANKRD22, high-affinity Fc-gamma receptor FCGR1 and apoptosis preventing NELL2.

**Fig 3.**
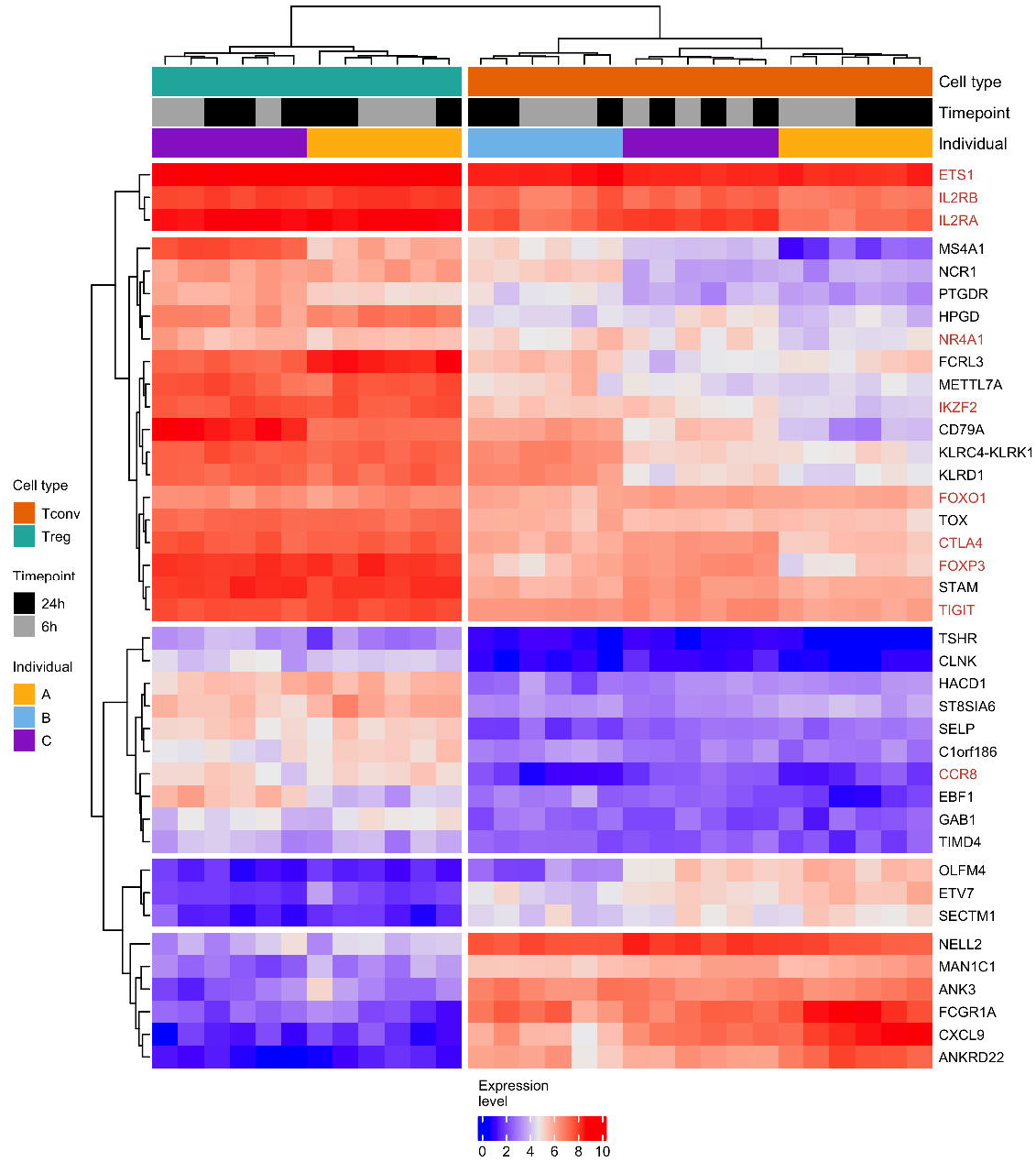
Gene expression differences between Tregs and Tconvs. The clustered heatmap shows log_2_ gene expression levels in Treg and Tconv samples included in the analysis. More precisely, the expression of top 30 differentially expressed genes together with the expression of differentially expressed Treg signature genes are shown. The annotations tracks show cell type, timepoint and individual whom the samples were obtained. Treg signature genes are coloured in red while other top differentially expressed genes are shown in black. Differential analysis was performed using Tregs from 2 males and Tconvs from 3 males from two timepoints. Individual was used as a covariate and p-values were extracted from linear regression models. The total number of tests was 11,453.

### Gene ontology analysis reveals T cell signalling and activation among genes having DMPs

A broad enrichment analysis showed that genes, where DMPs were mapped, were involved in T cell receptor signalling and T cell activation, leukocyte activation and differentiation, and various other general biological processes. In addition, many GO terms were associated with cellular migration and motility (**Fig S6**). The set of genes containing hypomethylated CpGs was enriched for the extracellular and ion transporter functions (**Fig S6**). We carried out similar analysis with genes containing DMR-s (**Fig S7**), however, it produced modest results. Next, we did a more detailed gene ontology analysis focusing on a collection C7 from ImmuneSigDB database, which contains immunologic signature genes deposited into Gene Expression Omnibus. We found significant enrichment among several studies focusing on CD4^+^ T cells e.g. Tregs, naïve, effector memory and central memory Th cells, adding further confidence that our analysis revealed methylation changes in T cell functional genes (**Fig S8**).

### Higher proportion of CpG sites with intermediate methylation levels in Tregs

We noted that the distribution of genome-wide methylation differed substantially between Tregs and Tconvs (**Fig 4**). While both had distinct bimodal distributions characteristic of cellular DNA methylation, Tconvs had a higher number of highly and lowly methylated (beta-value of 0.8-0.9 and 0-0.1, respectively) sites. In contrast, Tregs had more CpG sites with intermediate methylation (0.2-0.4) (**Fig 4A**). We found a similar change (**Fig S9**) in a published dataset [25]. More specifically, the CpG sites that had methylation levels of 0.2-0.85 in Tregs tended to have higher methylation in Tconvs (**Fig 4B**), indicating global hypomethylation in Tregs. In addition, we observed differences in the variability of methylation levels between Treg and Tconv cells. The methylation levels in Tregs were more variable compared to Tconvs (**Fig 4C**, Kolmogorov-Smirnov test’s p-value < 2×10^−16^) implying more diverse subpopulations. To see whether the global hypomethylation could be related to functional impact, we studied the methylation levels of CpG sites in hyper- and hypomethylated DMRs in Tregs and Tconvs. We saw that both hypo- and hypermethylated DMRs had a relatively similar distribution in Tregs while the distributions in Tconvs were vastly different. This indicated that in Tregs the methylation pattern shifts from two opposite modes into a middle area due to active regulation (Fig 4D-E) and suggested that not a site-specific transcription factor but a global epigenetic modifier should induce the methylation change in Tregs.

**Fig 4.**
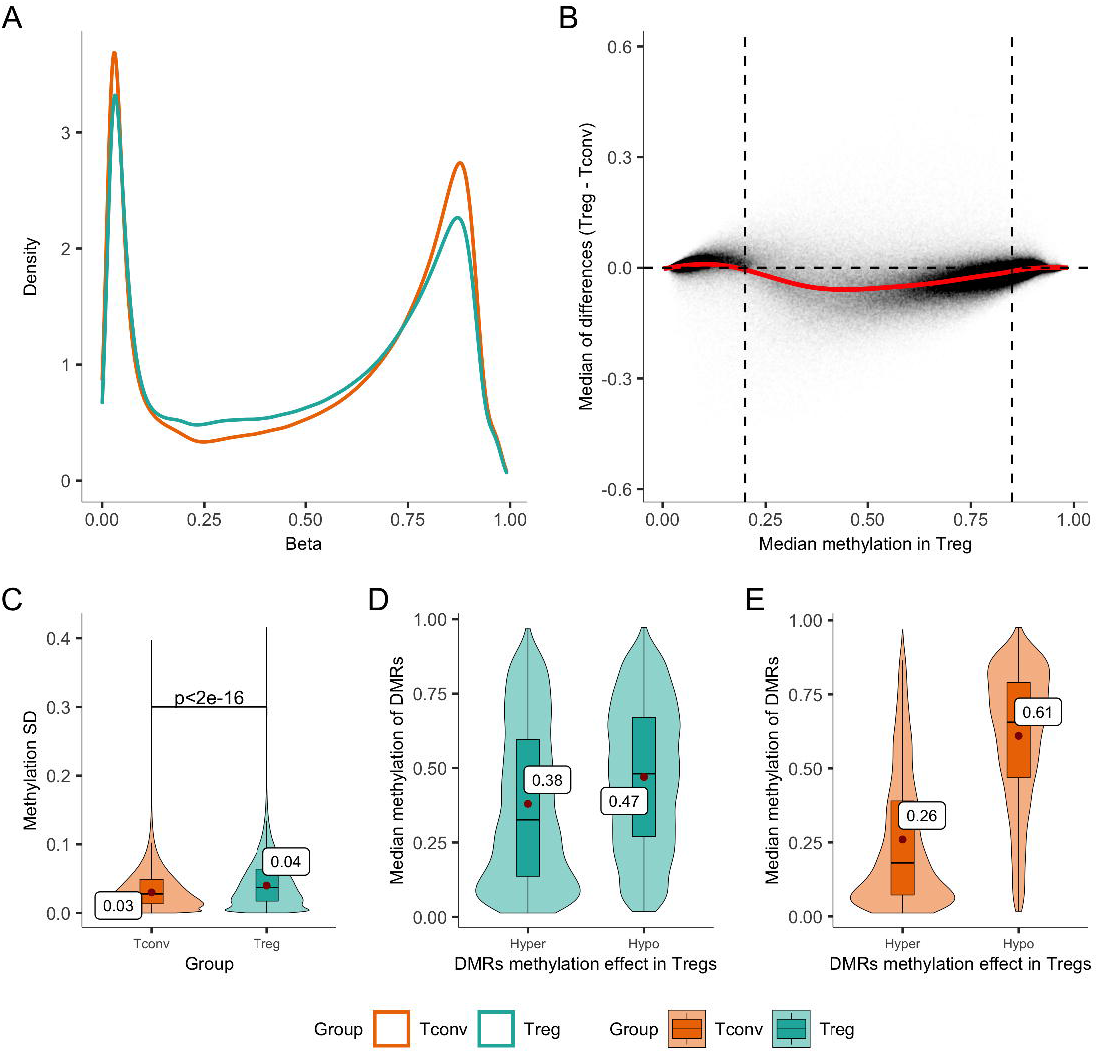
Global methylation in Tregs and Tconvs. (A) The density plot of methylation beta values across all studied sites (766,126) in Tconvs and Tregs. (B) Global methylation levels differ in Tregs. Treg median methylation levels are given on the x-axis and the difference of medians in Tconvs and Tregs on the y-axis. The horizontal line at 0 on the y-axis marks no change in medians and the red line corresponds to loess regression. The CpG sites with median methylation between 0.2 and 0.85 (the dotted vertical lines on the x-axis) tend to have hypomethylation in Tregs. (C) The higher degree of DNA methylation variance in Tregs is shown with boxplots. More specifically, each boxplot is overlaid with violin plot and represents the distribution of CpG wise calculated standard deviation (SD) of DNA methylation levels in Tregs and Tconvs. Two sample Kolmogorov-Smirnov test was used to obtain the p-value. (D-E) Boxplots showing median hyper- and hypomethylation of DMRs in Tregs and Tconvs, respectively. (A-E) The data is composed of DNA methylation profiles of Tregs and Tconvs from 6 individuals.

### Genes with differentially methylated sites contain Kaiso binding motifs

The changed distribution of global DNA methylation levels in Tregs in comparison to Tconvs suggested that an underlying transcriptional factor could affect the genome-wide methylation levels. The enrichment analysis of transcription factors using TRANSFAC data on gene sets containing hypo- and hypermethylated DMPs revealed enrichment of DNA methylation associated transcription factors Kaiso (ZBTB33) and Sp1. (Fig S6). Kaiso binds to methylated DNA [29], has been proposed to regulate DNA methylation homeostasis [30], and is expressed in Tregs, suggesting that it might function as one of the key regulators of Treg-specific demethylation. Its motif enrichment p-value in the hypo- and hypermethylated gene set is p=1.94×10^−13^ and p=1.18×10^−22^, respectively (Fig S6). In addition, multiple binding motifs of Sp1, another DNA methylation modifier, were highly enriched among both hypo- and hypermethylated CpG sites. Enrichment analysis also showed motifs of RNF96 (TRIM28), a part of the repressive chromatin remodelling complex, that is involved in transcriptional repression regulated by FOXP3 [31].

### DNA methylation changes in Tregs of GD patients

As we found the differential methylation of the TSHR gene in Tregs as compared to Tconvs, we studied the Treg-specific methylation in GD patients. Tregs proportions and FOXP3 expression levels are diminished in the GD patient peripheral blood samples [13, 14]. Considering the involvement of Tregs and the pathogenic role of TSHR in GD etiology, we investigated GD patients and healthy controls for their differential DNA methylation. We extracted Tregs from 11 GD patients and 10 controls and analysed their genomic DNA by Illumina EPIC methylation array. The DNA methylation differences between Tregs from GD patients and healthy individuals were relatively small and we observed no differences in the distribution of global methylation levels (**Fig S10**). The differential analysis resulted in only 19 differentially methylated sites with FDR ≤ 0.05 (**Sup. Table 7**). From these, 18 CpG sites were hypermethylated and 1 hypomethylated in Tregs from GD patients compared to healthy individuals (**Fig 5A**). Most of the DMP-s were located in the gene body and intergenic regions. Among several other CpGs implicated in autoimmune diseases (EP300, CBFA2T3, FMNL2, ADARB2 and PLCH2), the top DMP was in the promoter region of OPLAH, encoding 5-oxoprolinase, previously associated with a differentially expressed gene in a subset of SLE patients [32]. An additional DMP was present in SPAG6, expressed in primary and secondary lymphoid tissues with a role in immunological synapse formation. We also found 3 DVPs (**Sup. Table 9**), of which one was in SLC45A1, a proton-associated glucose transporter, and 2 DMRs (in RP11-420L9.4 and COX7A2L genes) (**Sup. Table 8**). However, we found no differential methylation at the CEP128/TSHR locus In GD patients (**Fig S11**).

**Fig 5.**
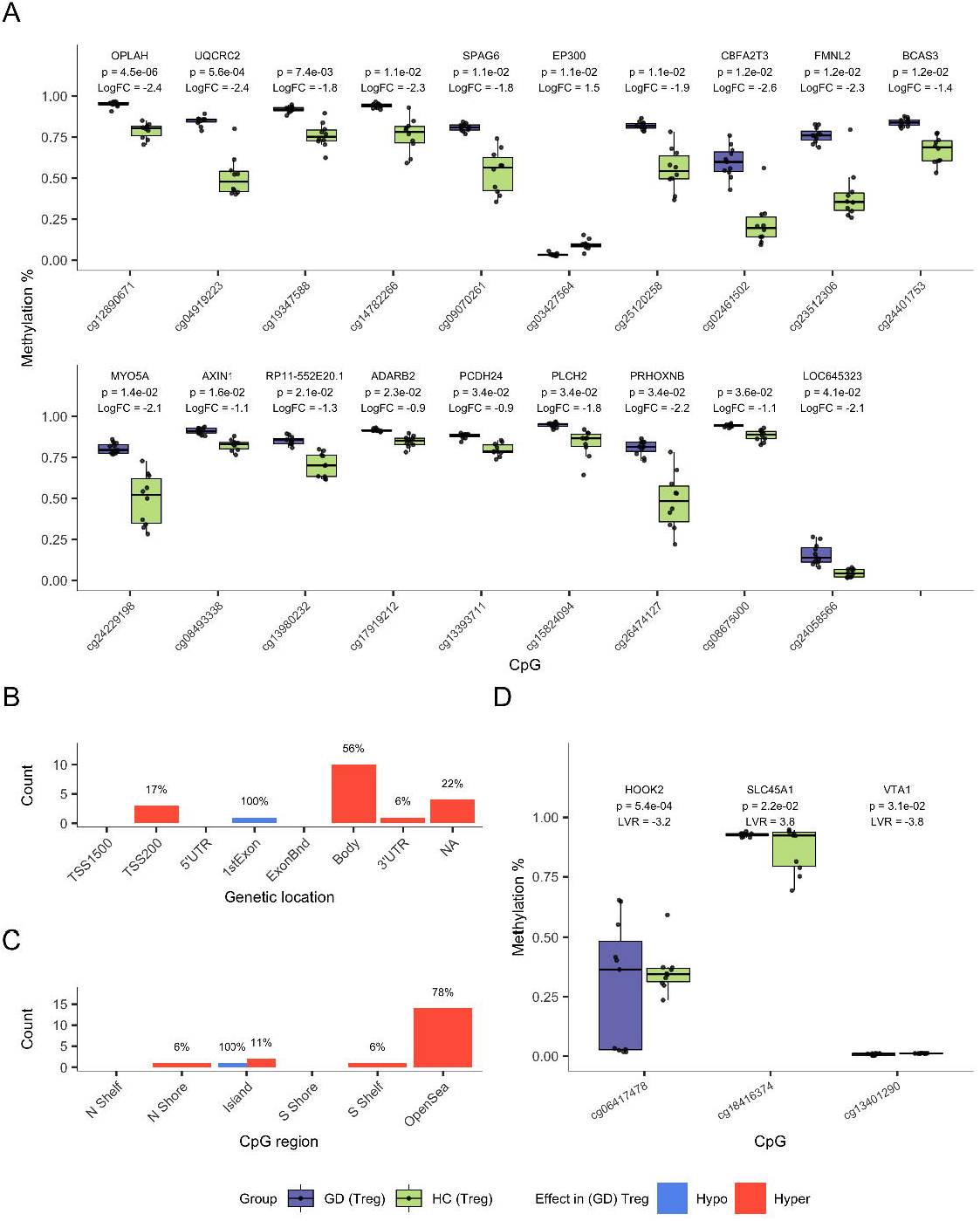
DNA methylation differences between Tregs from GD patients and healthy controls. (A) DMPs from differential analysis between Tregs of GD patients and healthy controls with the corresponding p-value (FDR-corrected), effect size (log_2_ fold change), and gene name (if nearby) (B-C) The location of DMPs in genes and CpG islands, respectively. The DMP count is shown relative to their location and percentages indicate hyper- and hypomethylated DMPs in a particular region. (D) DVPs at corresponding genes with FDR corrected p-value and effect size (log of variance ratio). (A, D) All p-values were obtained from linear regression models

## Discussion

Tregs and Tconvs are closely related cell types with opposing functional roles in their responses. Genome-wide studies revealed Tregs to have unique epigenetic modifications, which are acquired during their development [27, 33]. Our DNA methylation analysis in human Tregs identified over 16,000 differentially methylated CpGs in Tregs and Tconv. As previously reported, the majority of differentially methylated sites in Tregs were hypomethylated and preferentially located in gene bodies and not in promoter regions or in CpG islands. Among genes with top hypomethylated CpG sites, we found transcription factor *IKZF2* (Helios), a known marker of human Tregs [34]. Other top hypomethylated sites were in *CENPM*, coding inner protein of the kinetochore structure and *LYST*, a lysosomal trafficking regulator. The functional links of these two genes with Tregs remain unknown. One of the *CENPM* transcripts encodes a human minor leukocyte antigen with high expression in B cells [35] whereas *LYST* is involved in regulating the secretory processes of lysosomal vesicles and may affect the intracellular trafficking of CTLA-4 in T cells [36]. Similar to other studies, we found changed methylation at *TIGIT, IKZF4, CTLA4, BCL11B, IL2RA* and *TNFRSF9* genes with known functional roles in Tregs. The association of these CpG sites with active genes in Tregs suggests their location in regulatory regions for transcription. Among the hypermethylated regions, we also found an upstream region of Themis, which is one of the strongly downregulated genes by FOXP3 in response to TCR stimulation, suggesting that its suppression may be a part of the Treg developmental program [37, 38]. We also found the hypomethylation in intron 1 of FOXP3, close to the CNS1 and CNS2 regions. The hypomethylation of the FOXP3 intron 1 is needed for the generation of peripheral Tregs and the maintenance of FOXP3 expression in the thymic Tregs [3].

In comparison to Tconvs, we found an enhanced DNA methylation entropy in Tregs as we noted a shift from high or low methylation states to an intermediate fraction. The reason for this remains unknown, however, it is possible that the given effect arises due to increased diversities at Treg subpopulation levels or could indicate a global regulation by a certain set of transcription factors. Searching for the transcriptional regulators of putative DNA methylation we found a BTB/POZ subfamily member Kaiso, encoded by the ZBTB33 gene [29, 39]. Kaiso is a transcriptional repressor that is expressed in multiple cell types, including Tregs, and can affect the total genome methylation level by protecting gene regulatory regions from hypermethylation [30]. In this context, it is tempting to propose that the overall hypomethylation and lower stringency in methylation levels could be modulated in Tregs by Kaiso activity and its role in genomic DNA methylation homeostasis.

Interestingly, we found the TSHR gene and its neighbouring CEP128 gene locus hypomethylated in Tregs. TSHR has been associated in genome-wide association studies with GD and its protein is a well-established autoantigen in GD. Moreover, the polymorphism in the CEP128 was also identified as associated with GD without obvious linkage to the TSHR gene [28]. Because of the proximity of the two genes, the location of the CEP128 gene polymorphism in the enhancer region for TSHR cannot be ruled out. Furthermore, the Treg-specific transcriptome analysis showed the TSHR gene demethylation to be in concordance with its expression in Tregs.

The TSHR locus hypomethylation is probably specific to Tregs among other T cell populations. Our earlier analysis of bulk DNA methylation in CD4^+^ T cells showed hypermethylation of TSHR intron 1, which was increased in GD patients [40]. At least two different but not mutually exclusive scenarios have been proposed to explain the association of the TSHR gene with the loss of tolerance in GD [18–20]. The first proposes TSHR alternative splicing in the thyroid epithelial cells whereas the second is related to its differential expression in the thymus. We here propose a third, alternative scenario that the TSHR locus demethylation and upregulation in Tregs might have a role in immune tolerance to TSHR. To elucidate this, we analysed Treg DNA methylation profiles in healthy induvial and GD patient Treg populations. However, we found only 20 differentially methylated CpG sites and no differential methylation in the TSHR locus, indicating that the DNA methylation profiles in healthy and patient Tregs are similar. Despite being not differentially methylated between healthy controls and GD patients, the TSHR locus can be epigenetically different in specific Treg subtypes of GD patients, for example, in thymic Tregs, which we could not differentiate in our analysis.

In conclusion, we show that the TSHR gene is demethylated and upregulated in human Tregs and its putative functional role in Tregs and immune tolerance remains an intriguing topic for further studies.

## Data availability statement

The data underlying the statistical analysis of this article will be made available by the authors, without undue reservation.

## Authors’ contributions

PP and KK designed the experiments. AS performed the statistical analyses and prepared the figures under the guidance of SK and HP. LT, ML and KK performed the experiments and were responsible for the collection and documentation of the donor samples. AS and PP wrote the manuscript and KK, SK and HP made revisions to the manuscript. PP, LM and HP provided funding.

## Acknowledgments

We thank the participants of the study. The study was supported by The Centre of Excellence for Genomics and Translational Medicine funded by European Regional Development Fund (Project No.2014-2020.4.01.15-0012), and the Estonian Research Council grants PRG377, PRG1095 and PSG59.

