## Supplementary Figures for "Graves’ disease-associated TSHR gene is demethylated and expressed in human regulatory T cells"

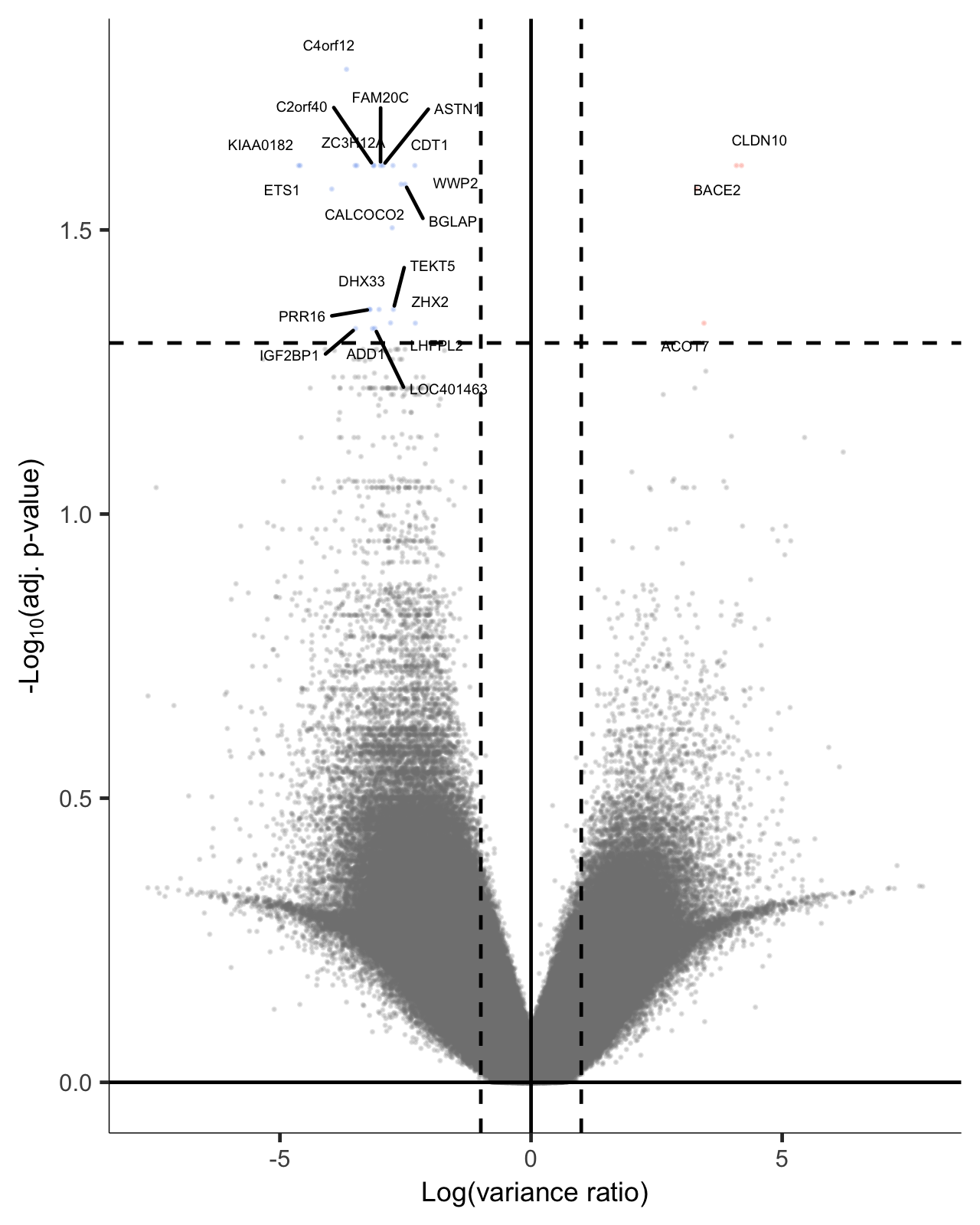

**Fig S1. Differential variability analysis of Tregs and Tconvs**. A volcano plot representing the results of differential variability analysis of Tregs and Tconvs from 6 individuals. Differentially variable CpG sites (DVPs) shown in blue indicate more variable and red less variable DVPs in Tregs. Additionally, the genes where top DVP-s reside are highlighted. The y-axis shows -log_10_ of FDR adjusted p-values and x-axis log_2_ of variance ratio. P-values were extracted from linear regression models and the total number of tests was 766,126 (the number of CpGs after pre-processing).

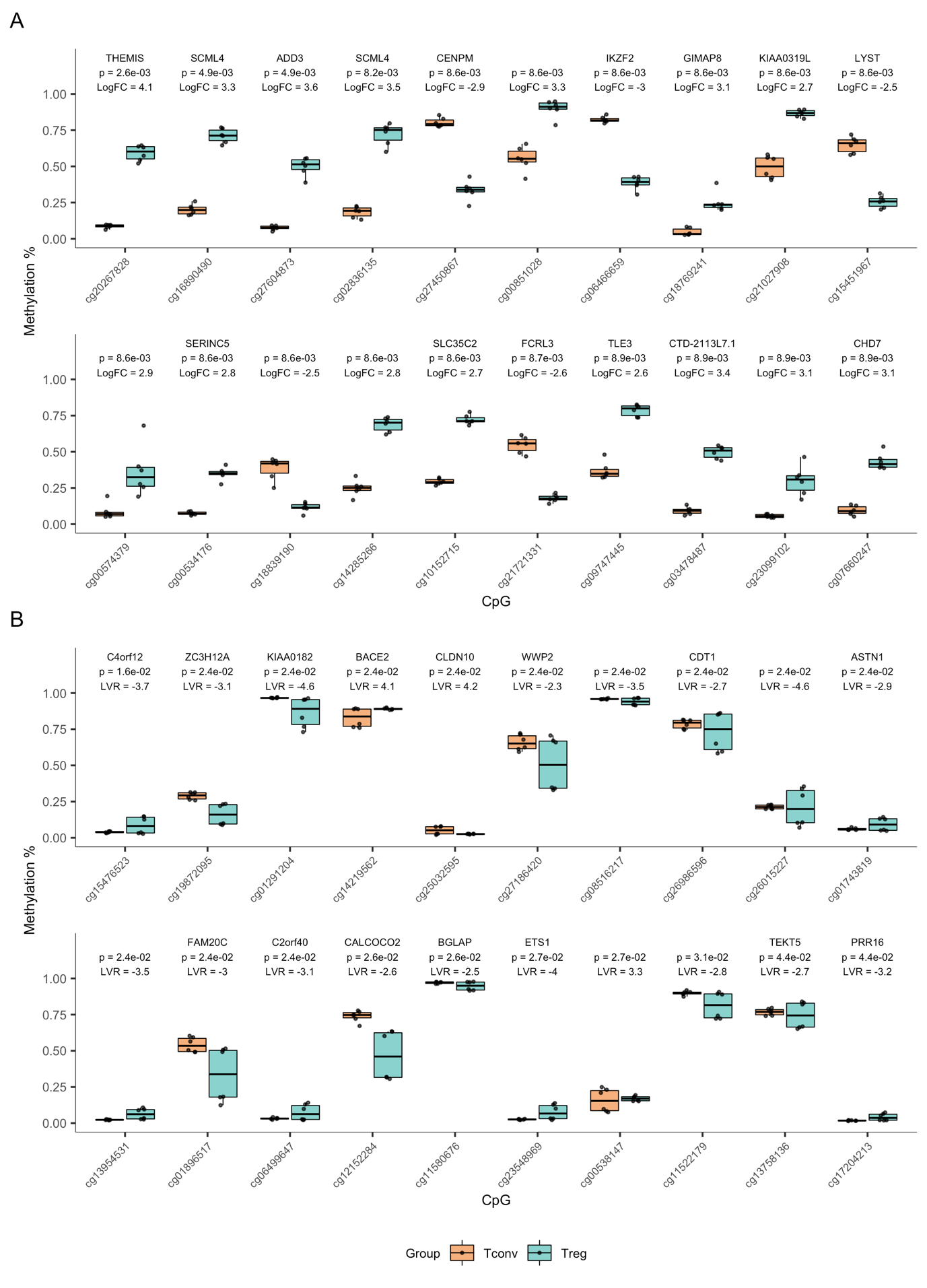
 **Fig S2. Top 20 DMPs and DVPs between Tregs and Tconvs.** Boxplots of top DMPs (A) and DVPs (B). The methylation levels of each CpG site are given with the corresponding FDR adjusted p-value, effect size and gene name (if nearby). LogFC stands for log fold change and LVR for log variance ratio. The differential analysis was carried out on Tregs and Tconvs from 6 individuals and p-values were extracted from linear regression models. The total number of tests was 766,126.

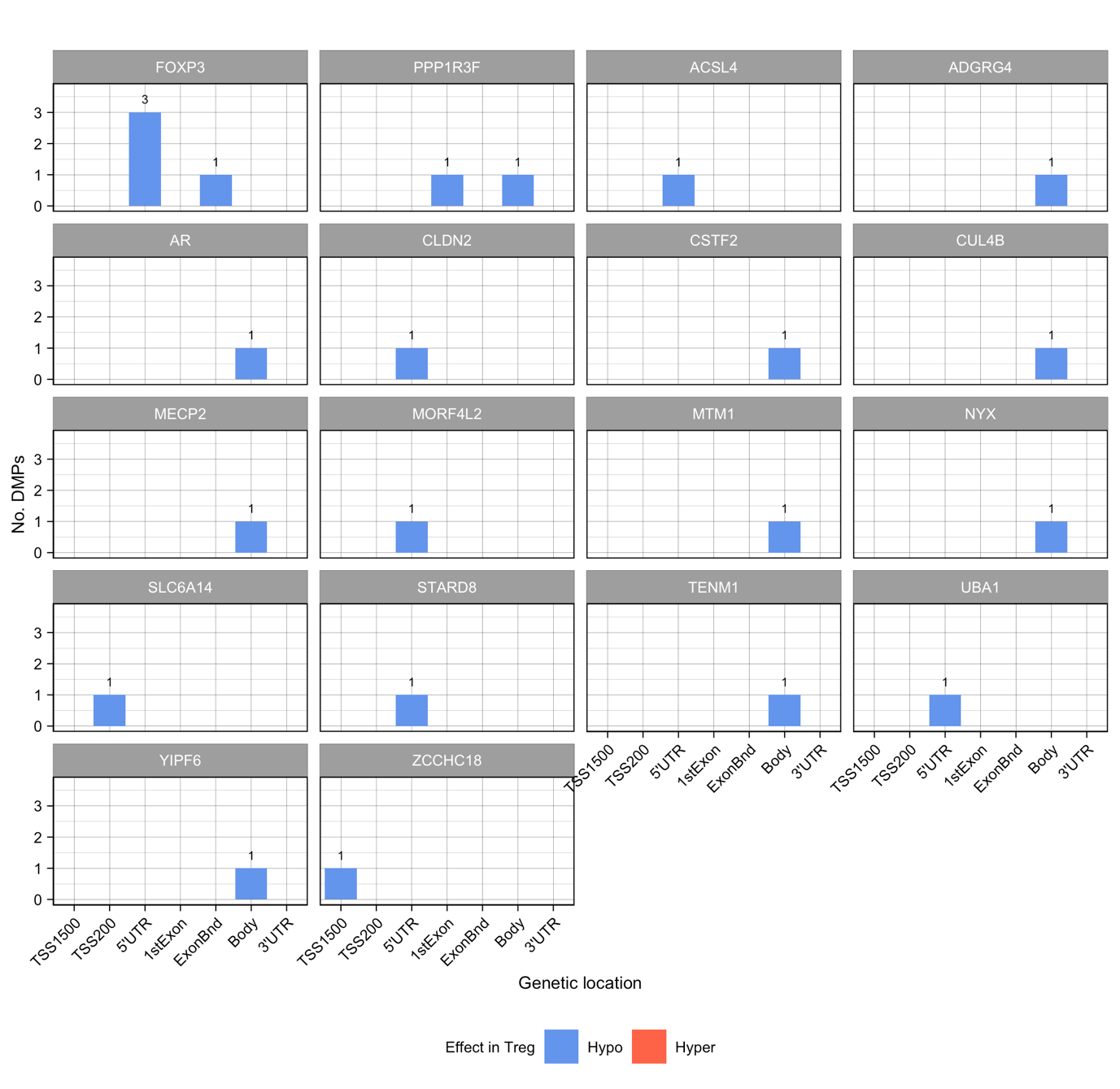

**Fig S3. Differential analysis of CpG sites on the X chromosome in Tregs and Tconvs.** The number of DMPs nearby each gene location. Colour marks the effect (change of methylation direction) in Tregs. Differential analysis was carried out on Tregs and Tconvs from 5 female participants and the total number of tests was 17,135. DMPs were determined based on FDR adjusted p-value (<0.05) from linear regression models where individual was included as a covariate.

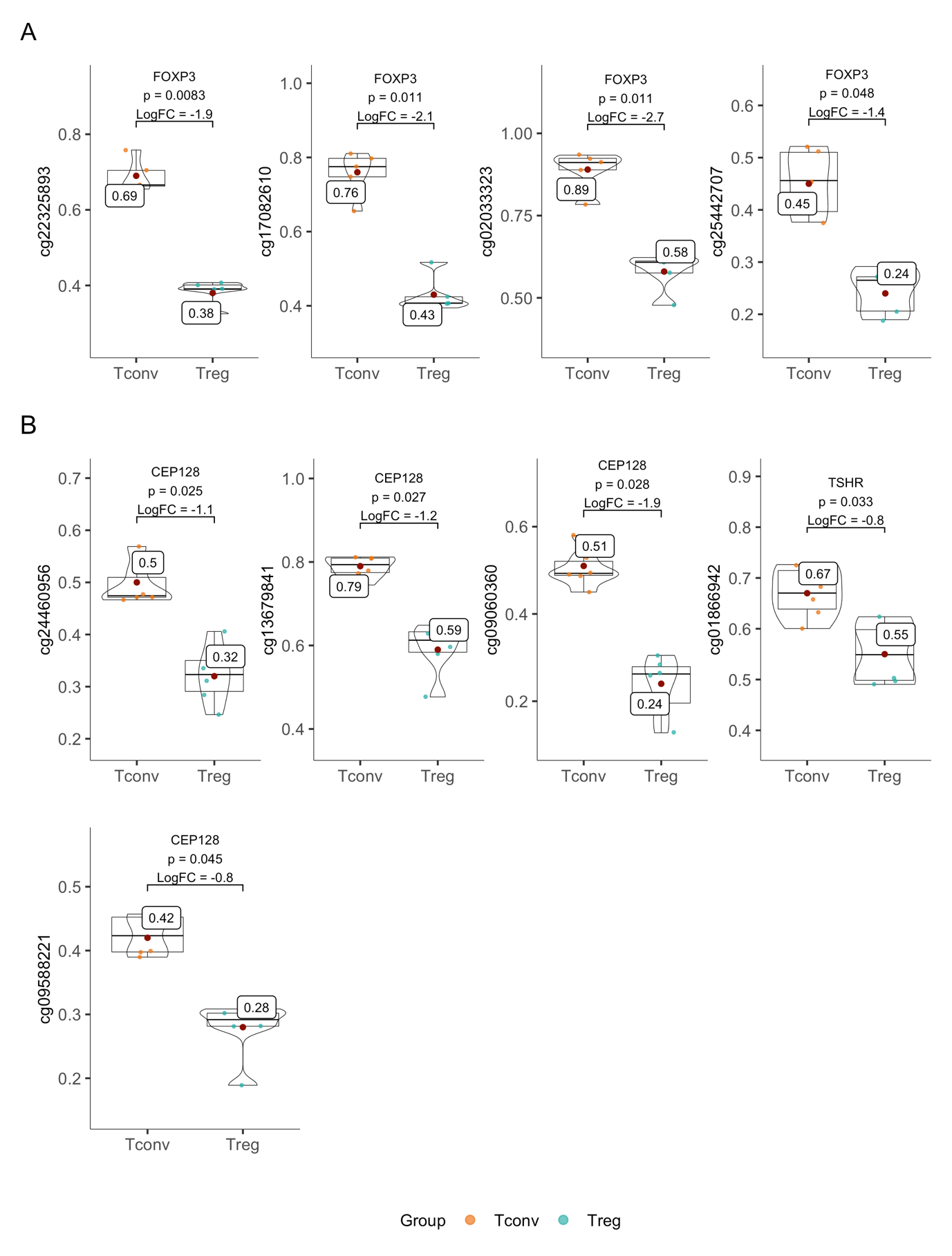

**Fig S4. Methylation levels of DMPs in FOXP3 and CEP128/TSHR loci**. Methylation levels of individual CpG sites are shown with boxplots overlaid with violin plots. Colour indicates the cell type, and each boxplot is annotated with gene name, FDR adjusted p-value, logFC (log fold change) and mean methylation level. P-values were obtained from linear regression based differential analysis between Tregs and Tconvs. Group sizes were 5 for FOXP3 (X chromosome was analysed separately using Tregs and Tconvs from 5 female participants) and 6 for CEP128/TSHR loci.

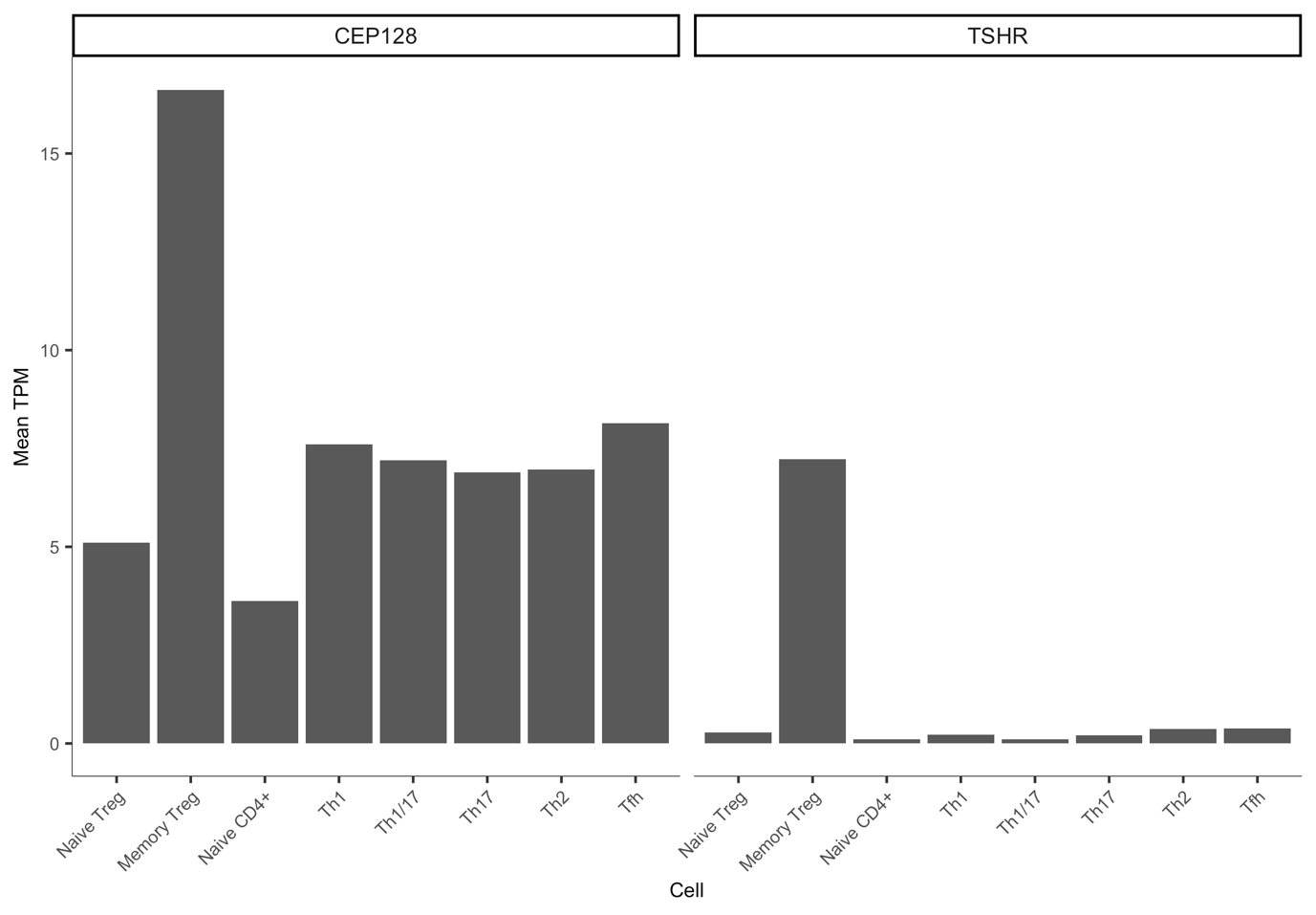
 **Fig S5. CEP128 and TSHR expression in CD4^+^ T cells in DICE database**. Mean TPM (transcripts per million) over all samples is shown for each CD4^+^ T cell subset.

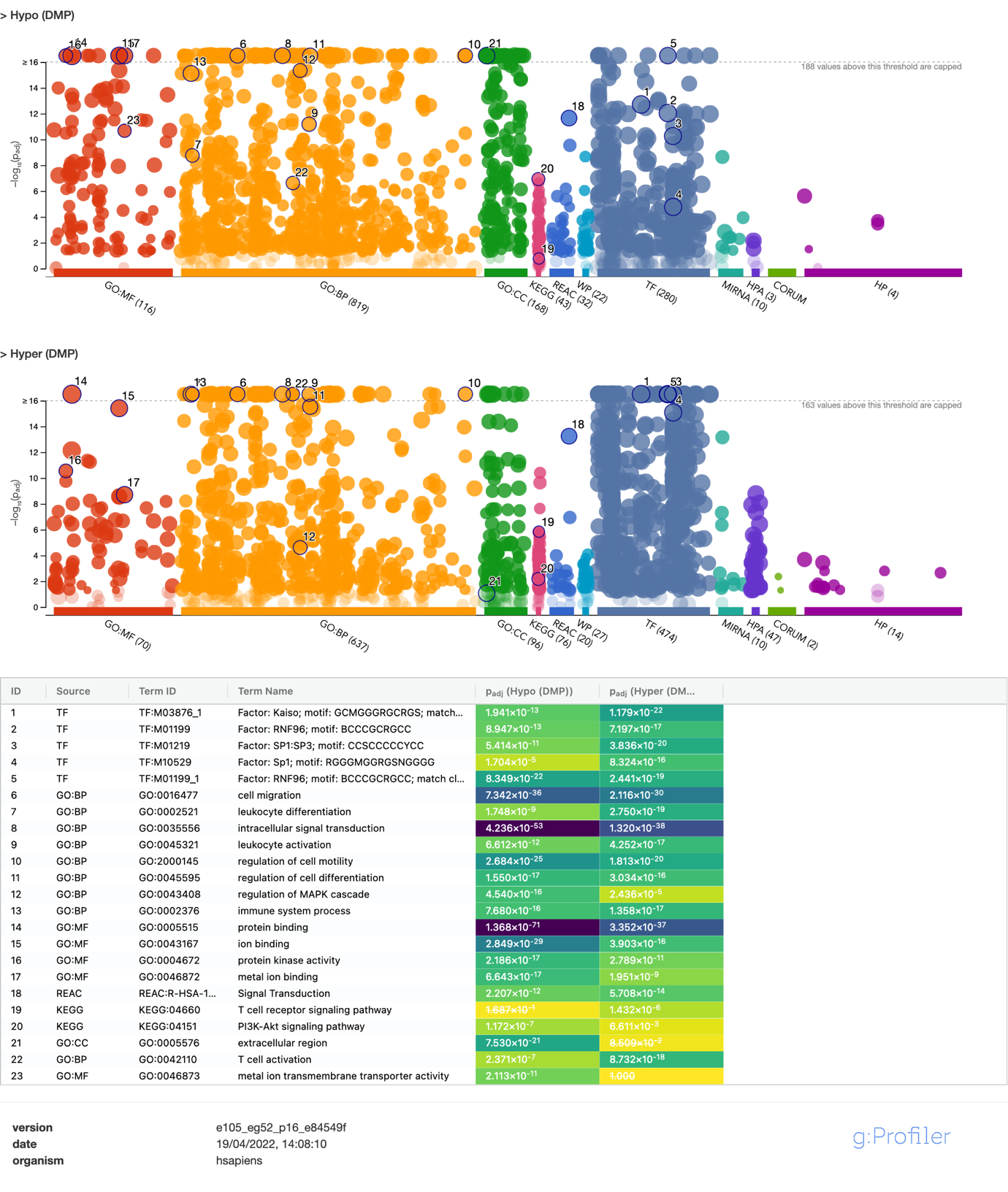

**Fig S6. Gene enrichment analysis of genes with DMPs**. Manhattan plots of enrichment analysis results done with g:Profiler web service [24]. The results of two gene sets are shown with upper plot corresponding to the gene set consisting of genes that contained hypomethylated DMPs and lower to genes that contained hypermethylated DMPs. Methylation effect is given in respect to Tregs (e.g., hypomethylated DMP has lower methylation level in Tregs compared to Tconvs). The y-axis of those plots corresponds to -log_10_ of g:SCS adjusted p-values from hypergeometric tests and x-axis corresponds to ontology terms that are grouped by their ontologies and subontologies. In addition, a table of selected ontology terms is shown. Link to results: <https://biit.cs.ut.ee/gplink/l/QaP_DFNaR5>

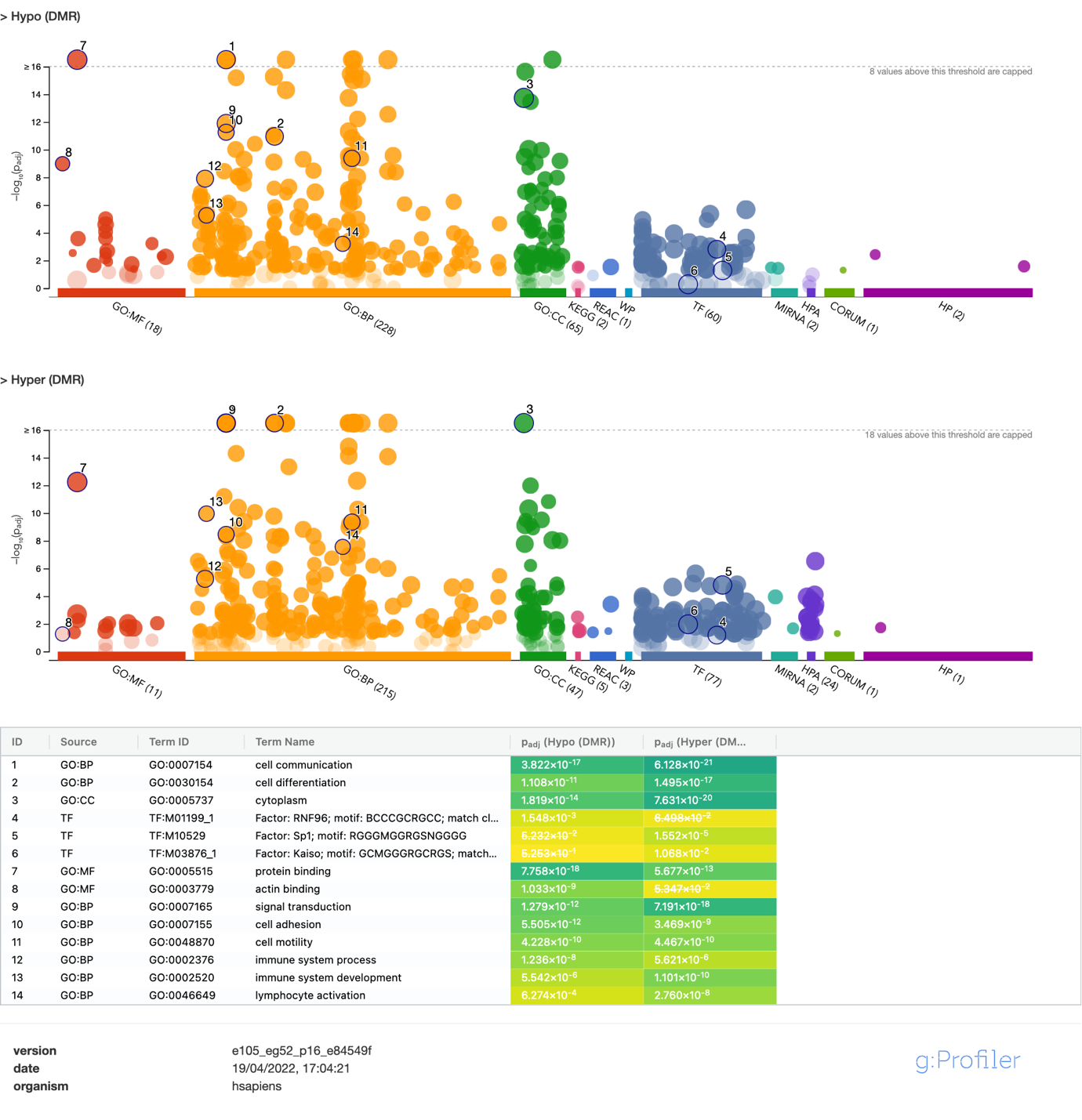
 **Fig S7. Gene enrichment analysis of genes with DMRs**. Enrichment analysis results of genes containing DMRs done using g:Profiler web service [24]. Upper Manhattan plot shows results of genes containing hypomethylated DMRs and lower hypermethylated DMRs. Methylation effect indicates the methylation direction in relative to Tregs (e.g., hypomethylated DMR has on average lower methylation level in Tregs compared to Tconvs over all CpGs sites belonging to a given DMR). The y-axis of those plots corresponds to -log_10_ of g:SCS adjusted p-values from hypergeometric tests and x-axis corresponds to ontology terms that are grouped by their ontologies and subontologies. Additionally, a table of selected ontology terms is shown. Link to results: <https://biit.cs.ut.ee/gplink/l/M67LVDb7SY>

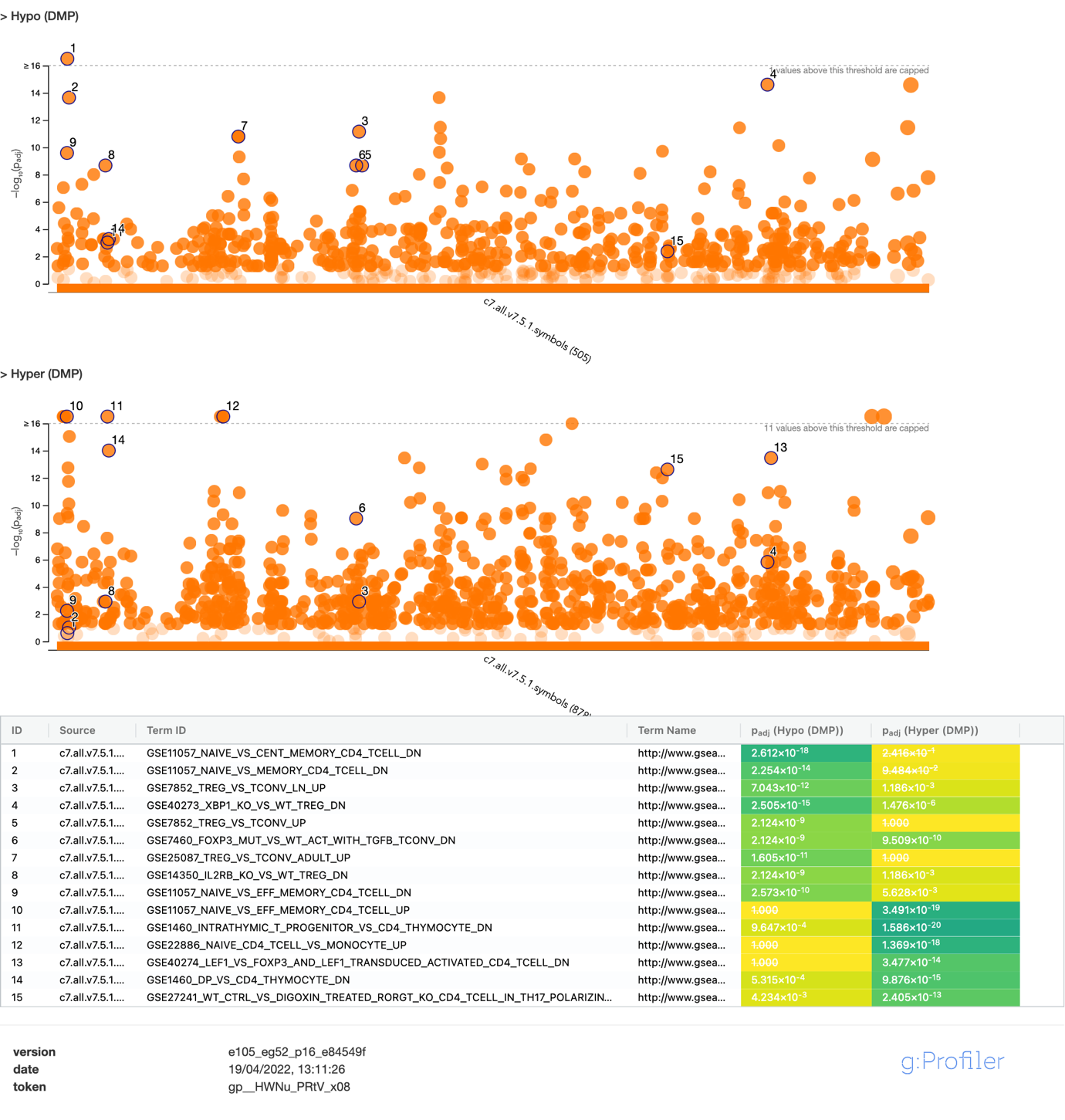

**Fig S8. Enrichment analysis of genes with immunological annotation data**. Similarly to Fig S5, given figure shows results of enrichment analysis of genes where DMPs mapped, however, in this case alternate annotation data was used. The y-axis of those plots corresponds to -log_10_ of g:SCS corrected p-values from hypergeometric tests and x-axis corresponds to ontology terms from MSigDB (Molecular Signatures database) C7 collection that contains immunologic signature gene sets. In addition, a table containing selected ontology term information is shown. Link to results: <https://biit.cs.ut.ee/gplink/l/rdVJJTKBSq>

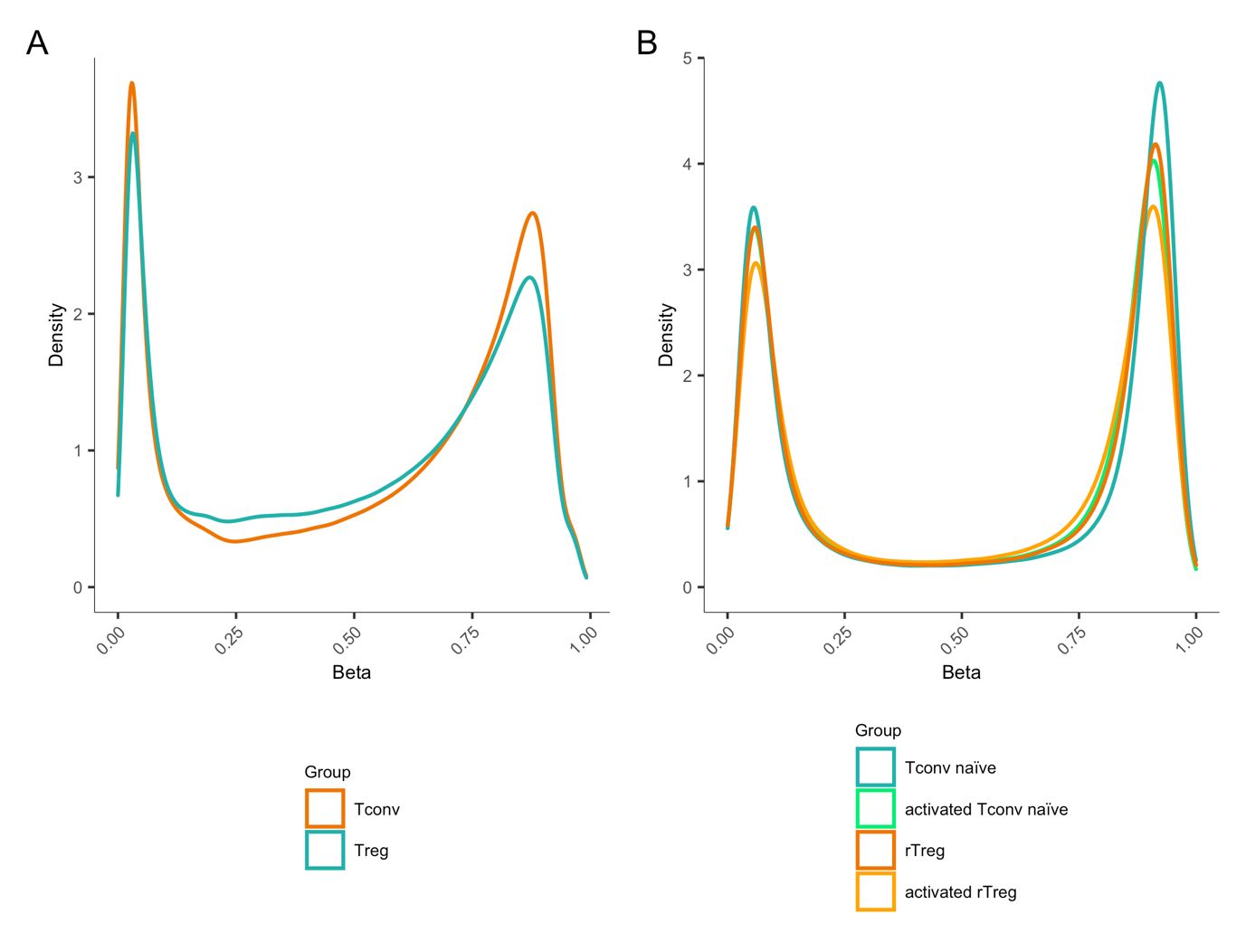

**Fig S9. Distribution of DNA methylation levels in Treg and Tconv cells.** Density plots of methylation levels in Treg and Tconv cells in our (A) and Zhang et al dataset [25] (B). The plots show that Tconvs’ bimodal methylation distribution is more enriched in high and low methylation values in comparison to Tregs.

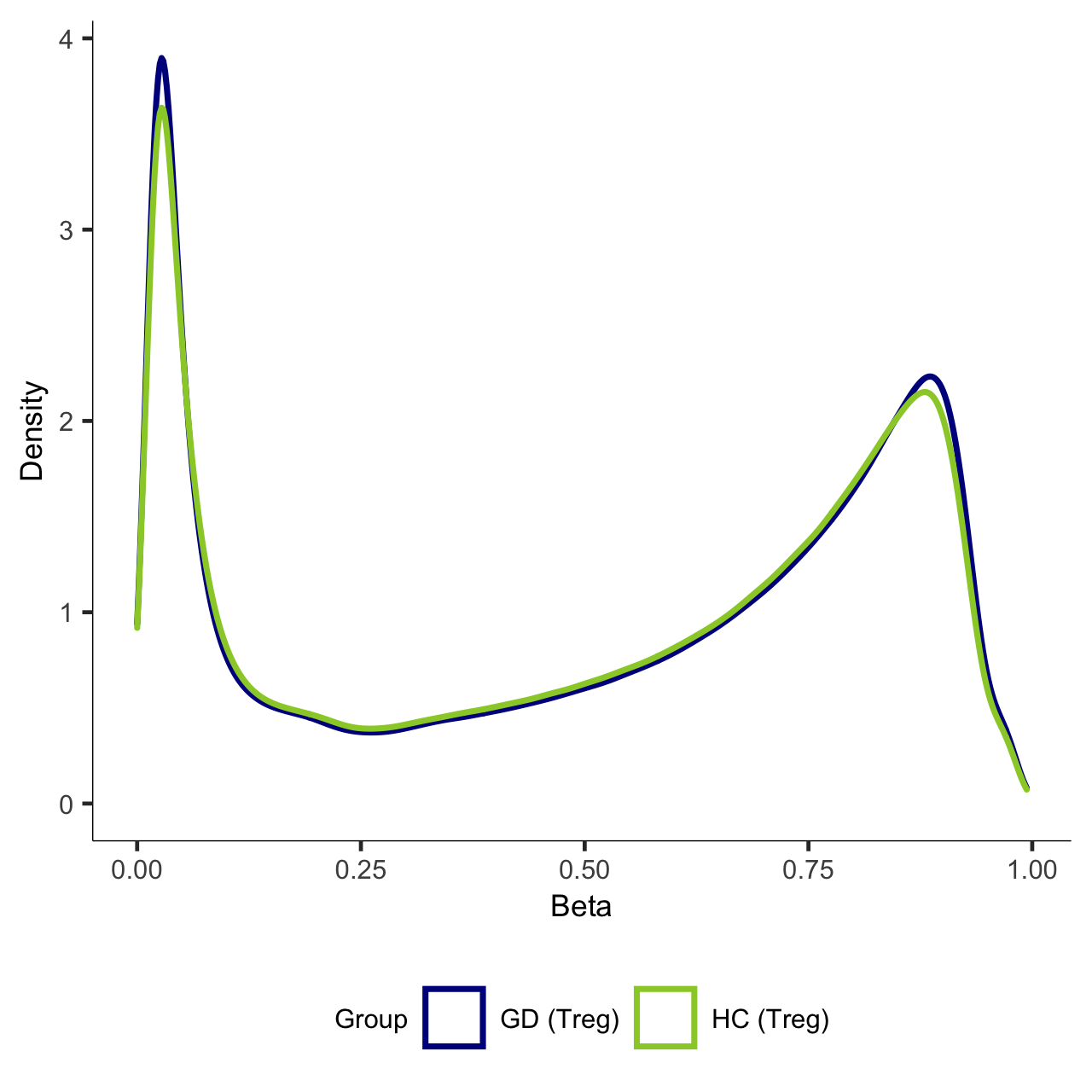

**Fig S10**. **Distribution of global methylation levels in Tregs from Graves’ patients and healthy controls.**

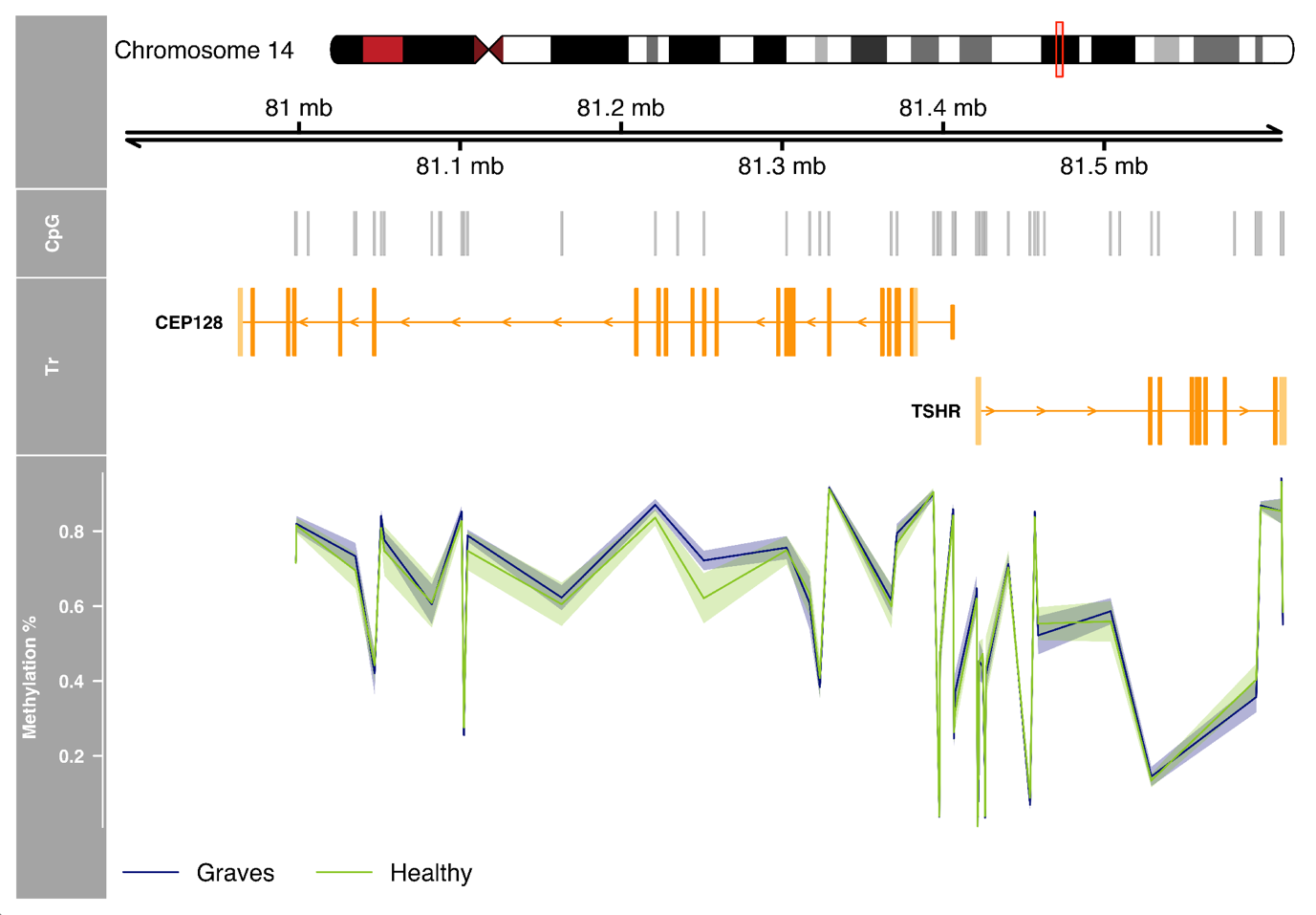

**Fig S11.** **Methylation levels of CpG sites in CEP128/TSHR regions in Tregs from Graves’ patients and healthy controls together with transcript and chromosomal information.**
