## Supplementary Table 10 for "Graves’ disease-associated TSHR gene is demethylated and expressed in human regulatory T cells"

Study	Sample	Group size	Sex (F/M)	Age range (ave±SD)

Treg vs Tconv

(DNA methylation)

Tconv	6	(5/1)	24-60 (40.5±14.6)

Treg	6	(5/1)	24-60 (38.5±14.2)

GD vs healthy (Tregs)

(DNA methylation)

GD	11	(3/8)	22-62 (44.8±13.2)

Healthy	10	(3/7)	25-61 (43.6±11.7)

Treg vs Tconv

(gene expression)

Tconv	3	(0/3)	33-48 (40.3±7.5)

Treg	2	(0/2)	40-48 (38.5±14.2)
